# Botulinum neurotoxin A modulates the axonal release of pathological tau in hippocampal neurons

**DOI:** 10.1101/2023.02.13.528198

**Authors:** Chiara Panzi, Sunaina Surana, Samantha De La-Rocque, Edoardo Moretto, Oscar Marcelo Lazo, Giampietro Schiavo

**Author notes:** Corresponding author: Department of Neuromuscular Diseases, Queen Square Institute of Neurology, Queen Square, WC1N 3BG (UK). Email: Chiara Panzi, Giampietro Schiavo.

## Abstract

Pathological tau aggregates propagate across functionally connected neuronal networks in human neurodegenerative pathologies, such as Alzheimer’s disease. However, the mechanism underlying this process is poorly understood. Several studies have showed that tau release is dependent on neuronal activity and that pathological tau is found in the extracellular space in free form, as well as in the lumen of extracellular vesicles. We recently showed that metabotropic glutamate receptor activity and the SNAP25 integrity modulate the release of pathological tau from human and mouse synaptosomes. Here, we have leveraged botulinum neurotoxins (BoNTs), which impair neurotransmitter release by cleaving specific synaptic SNARE proteins, to dissect molecular mechanisms related to tau release at synapses. In particular, we have tested the effect of botulinum neurotoxin A (BoNT/A) on the synaptic release of tau in primary mouse neurons. Hippocampal neurons were grown in microfluidic chambers and transduced with lentiviruses expressing human tau (hTau). We found that neuronal stimulation significantly increases the release of mutant hTau, whereas wild-type hTau is unaffected. Importantly, BoNT/A blocks mutant hTau release, indicating that this process is modulated by SNAP25 in intact neurons. These results suggest that BoNTs are potent tools to study the spreading of pathological proteins in neurodegenerative diseases and will play a central role in identifying novel molecular targets for the development of therapeutic interventions to treat tauopathies.

## Introduction

Tauopathies are a group of neurodegenerative diseases characterised by microtubule-associated protein tau (MAPT) aggregation. They include Alzheimer’s disease (AD; (Braak & Braak, 1991; Cope et al., 2018; Sanders et al., 2014; Scholl et al., 2016; Tracy et al., 2022), Pick’s disease (PiD), corticobasal degeneration (CBD), frontotemporal dementia (FTD) and several other rarer diseases (Goedert & Spillantini, 2019; Virginia M-Y Lee et al., 2001). The discovery of the involvement of MAPT mutations in FTD associated with parkinsonism (FTDP; (Hutton et al., 1998; Poorkaj et al., 1998; Spillantini et al., 1998) and several other pathologies (Goedert & Spillantini, 2019), together with the lack of success in clinical trials focused on amyloid beta (Aβ) pathology in AD, highlighted the importance of studying tau as a potential target for the treatment of neurodegenerative diseases associated with dementia.

Tau is encoded by the *MAPT* gene located on human chromosome 17q21.31. The alternative splicing of its mRNA generates six different isoforms characterised by the presence of three or four C-terminal tandem repeats (3R or 4R) and of zero, one or two repeats (0N, 1N or 2N) at the N-terminus (Goedert & Spillantini, 2019; Virginia M-Y Lee et al., 2001). Tau 1N4R is the most expressed isoform in the adult human brain (Hong et al., 1998) and, like the other 4R isoforms, it can harbour mutations that are found in familial tauopathies, as P301L and P301S (Goedert & Spillantini, 2019).

Tau aggregates forming intracellular neurofibrillary tangles (NFTs), which progressively accumulate in the central nervous system ((Cipriani et al., 2011), and it propagates along synaptically connected regions (Braak & Braak, 1991; Hoenig et al., 2018; Vogel et al., 2020) in a well-defined, staged pattern that correlates with the cognitive decline seen in AD patients (Braak et al., 2006). *In vivo* and *in vitro* studies have shown that pathological protein aggregates, such as α-synuclein and hyperphosphorylated tau, spread in connected neural networks, similarly to the prion protein (Brundin et al., 2010; Walker & Jucker, 2015). Emerging evidence also suggests that this spreading of pathological proteins in neuronal-connected regions is activity-dependent (Pooler et al., 2013; Wu et al., 2016; Yamada et al., 2014).

Although tau is associated with different organelles, including ectosomes (Dujardin et al., 2014), exosomes (Wang et al., 2017) and synaptic vesicles (McInnes et al., 2018), and tau oligomers are transported along axons and dendrites (J. W. Wu et al., 2013), the precise mechanisms controlling tau release and reuptake in neurons are not completely understood (Xu, 2021). Several distinct mechanisms have been suggested for tau secretion, such as active exocytosis and unconventional secretion pathways (Katsinelos et al., 2021; Katsinelos et al., 2018; Nickel & Rabouille, 2009). Extracellular tau has been found both inside extracellular vesicles (EVs; (Gabrielli et al., 2022), comprising exosomes, ectosomes and apoptotic bodies, and in its free form (Mathieu et al., 2019). Free tau represents the majority of extracellular tau (90%), whereas just 7% and 3% is found in ectosomes and exosomes, respectively (Dujardin et al., 2014). Although the majority of extracellular tau is in its free form, it is not clear which tau species is responsible for interneuronal spread in tauopathies. Therefore, tau in EVs could contribute to the overall pathogenic process (De La-Rocque et al., 2021; Gabrielli et al., 2022). Indeed, several lines of evidence support the role of exosomes, single membrane vesicles with a diameter between 50 and 150 nm (Mathieu et al., 2019). Exosomes are generated by the direct fusion of multivesicular bodies (MVBs) with the plasma membrane (Mathivanan et al., 2010). Synaptic activity regulates the release of exosomes (Bahrini et al., 2015) as well as other neuronal secretory organelles (Padamsey et al., 2017), representing an important pathway for intercellular communication (Pegtel & Gould, 2019).

The association of tau with exosomes (Wang et al., 2017), synaptic vesicles (McInnes et al., 2018; Zhou et al., 2017) as well as the soluble N-ethylmaleimide-sensitive factor attachment protein receptor (SNARE) proteins (Lee et al., 2021; Pilliod et al., 2020; Tracy et al., 2022; Xu, 2021) suggests that the neuronal spreading of tau may be dependent on the integrity of synaptic SNAREs, such as syntaxin, 25 kDa synaptosomal-associated protein (SNAP25) and vesicle-associated membrane protein (VAMP)/synaptobrevin. Interestingly, these proteins can be selectively impaired by clostridial neurotoxin (CNT) family, which comprises tetanus neurotoxin (TeNT) and several botulinum neurotoxins (BoNTs). CNTs block neuroexocytosis by cleaving distinct SNARE proteins at different sites. Specifically, TeNT and BoNT/B, /D, /F, /G and /X cleave VAMP/synaptobrevin, BoNT/A and /E cleave SNAP25, whereas BoNT/C cleaves both syntaxin and SNAP25 (Rossetto et al., 2021; Schiavo et al., 2000). As a result, the proteolytic activity of CNTs can be detected in cells and tissues with antibodies specific for SNARE epitopes generated by the action of these neurotoxins in treated neurons (Caleo et al., 2018; Fabris et al., 2022).

TeNT and BoNTs are both synthesised as a single-chain protein of 150 kDa, which is cleaved by proteases into a 50 kDa light chain (L) and a 100 kDa heavy chain (H). The two chains are linked via a disulphide bridge, which is essential for neurotoxicity (Schiavo et al., 1990). Three different functional domains are required for neuronal entry and activity of CNTs. The L chain contains the Zn^2+^-dependent metalloprotease activity; the N-terminal part of the H chain (H_N_) is required for pore formation and L chain translocation into the cytosol, whilst the C-terminal part of the H chain (H_C_), is responsible for neuronal binding (Schiavo et al., 2000; Montal, 2010).

The ability of BoNTs to block neurotransmitter release, by inhibiting synaptic vesicle fusion with the plasma membrane, makes them ideal tools to study secretory events in mammalian neurons (Mazzo et al., 2022; Montal, 2010). Furthermore, BoNTs are widely used in clinic because of their potency, neuronal specificity, and reversibility (Pirazzini et al., 2017). These features make them preferred therapeutics for the treatment of several human conditions characterised by hyperactivity of nerve terminals, such as spasticity, dystonia, and hyperactive bladder (Pirazzini et al., 2017). Some of the therapeutic effects of BoNTs are due to their activity at distal loci following axonal transport (Restani, Giribaldi, et al., 2012). Whilst BoNTs show a preference for cholinergic terminals, they can also affect other neuronal endings, such as GABAergic and glutamatergic synapses (Caleo et al., 2018). This suggests that the *in vivo* use of BoNTs may be expanded to the study and treatment of pathologies characterised by hyperactivity of non-cholinergic neurons.

Recent studies have exploited BoNTs as tools to study molecular mechanisms regulating the secretion of pathological aggregates (Mazzo et al., 2022; Okuzumi et al., 2018). In particular, it was shown that the spreading of α-synuclein aggregates injected in the mouse brain decreases after treatment with BoNT/B in the contralateral hemisphere (Okuzumi et al., 2018). Mazzo et al. demonstrated that cleavage of SNAP25 by BoNT/A or sequestration of SNAP25 by a metabotropic glutamate receptor agonist modulate the release of pathologic tau from mouse and human synaptosomes (Mazzo et al., 2022). Altogether, these studies show the modulation of hTau release by specific SNARE proteins, suggesting that pathological hTau secretion might be dependent, at least partially, on the membrane fusion machinery controlled by synaptic SNAREs.

In this work, we exploit the activity of BoNT/A to elucidate the molecular mechanisms regulating the spreading of hTau aggregates across neuronal circuits. Using this approach, we aim to provide novel insights into the mechanisms controlling pathological hTau release from synaptic terminals and identify novel molecular targets for the development of therapeutic interventions to treat tauopathies.

## Results

### Clostridial neurotoxin activity in primary neurons

We first wanted to determine the concentration at which the commercial preparation of BoNT/A Dysport was active in our system and if it was cleaving SNAP25 as expected. We converted the units (U) of toxins in molar concentration using the amount of active BoNT/A found in Dysport (0.87 ng/100 U; (Pirazzini et al., 2017). We then tested BoNT/A activity in primary hippocampal neurons in the absence of external neuronal stimulation. At 14 days in vitro (DIV14), primary hippocampal neurons were treated with increasing concentrations (5 pM, 8.7 pM, 17.3 pM) of BoNT/A for 24 h in 250 μl of neuronal medium. The maximum volume of toxin added was set to 10% of the total media volume to minimise toxicity, and BoNT/A activity was determined by immunohistochemistry using an antibody specifically recognising BoNT/A-cleaved SNAP25 (cl.SNAP25, **Figure 1B**) (Fabris et al., 2022). This antibody does not recognise full-length SNAP25, as shown by the negligible background signal in untreated cells (**Supplementary Figure 1**). BoNT/A activity was measured in synapses (synaptophysin mask, **Figure 1B-C**) and at the whole neuron level (βIII tubulin mask, **Figure 1B-C**) to determine the efficiency of BoNT/A in cleaving SNAP25. Consistent with the broad neuronal distribution of SNAP25 (Hagiwara et al., 2005; Tao-Cheng et al., 2000), we detected cl.SNAP25 not only at synapses, but also along neurites and in the soma, as indicated by the similar intensity of cl.SNAP25 we measured in both the synaptophysin and βIII tubulin domains (**Figure 1C**).

**Figure 1.**
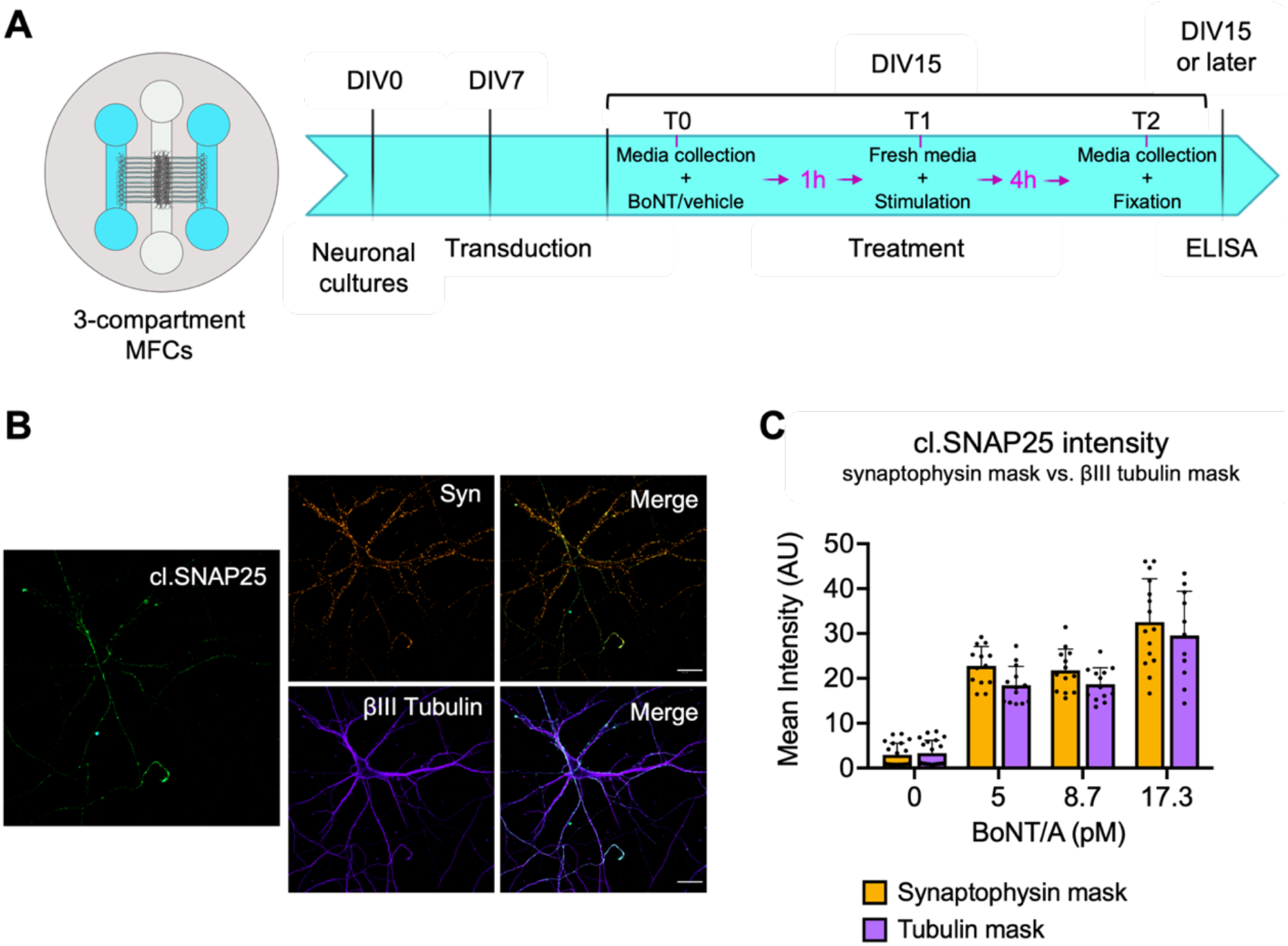
Experimental set-up for hTau release in microfluidic chambers and BoNT/A activity in vitro. A) Schematic of the 3-compartment microfluidic chambers (MFC) used in our experimental set up (left panel). Timeline for measuring hTau release in primary hippocampal neurons in MFCs (right panel). The numbers of days in vitro (DIV) when primary hippocampal neuron cultures were treated is shown. At DIV15, neurons were sequentially treated as follows: at time point 0 (T0), media was collected and BoNT/A (Dysport) or 0.9% NaCl (vehicle) was added in the axonal compartment; at time point 1 (T1, 1 h incubation), neurons were stimulated and at time point 2 (T2, 4 h incubation), media was collected and neurons were fixed. B) Representative images of primary hippocampal neurons treated with 17.3 pM of Dysport for 24 h. Activity is determined by measuring the signal of cleaved SNAP25 (cl.SNAP25, green) in the synaptophysin (Syn, orange) and βIII tubulin (purple) positive areas. Scale bar = 25 μm. C) BoNT/A activity is quantified based on the anti-cl.SNAP25 antibody signal intensity. The BoNT/A activity in the two different masks (synaptophysin mask vs. tubulin mask) is shown. A two-way ANOVA with Šídák’s multiple comparisons test was performed, and all possible pairs were compared. There is no significant difference between the two masks. n = 3. Panel A was created using Biorender.com.

### Expression of 1N4R hTau using a lentiviral vector

Of the six human tau isoforms generated by alternative splicing of the *MAPT* gene, the 1N4R isoform is the most expressed in the adult brain (Hong et al., 1998). Several mutations in the 4R tau isoforms, such as the P301S mutation used in this study, are found in familial tauopathies (Goedert & Spillantini, 2019; Virginia M-Y Lee et al., 2001). In tauopathies, tau forms pathologic aggregates that are able to spread across connected neuronal networks (Vogel et al., 2020; Vogels et al., 2020). In this project, we aimed to set up a model system to determine the effect of tau P301S familial mutation on tau spread. Therefore, we generated viral vectors encoding the 1N4R hTau isoform, either wild-type (WT) or harbouring the P301S mutation (P301S). Each of these was tagged with a FLAG sequence at the carboxy terminus to enable discrimination between overexpressed hTau and endogenous murine tau (**Figure 2A**). Robust expression of hTau by the plasmid (**Figure 2B**) as well as the derived lentivirus (**Figure 2C**) was detected in DIV15 primary hippocampal neurons. When cultured in microfluidic chambers (MFCs), primary neurons expressed hTau in both the somatic and axonal compartments (**Figure 2C**).

**Figure 2.**
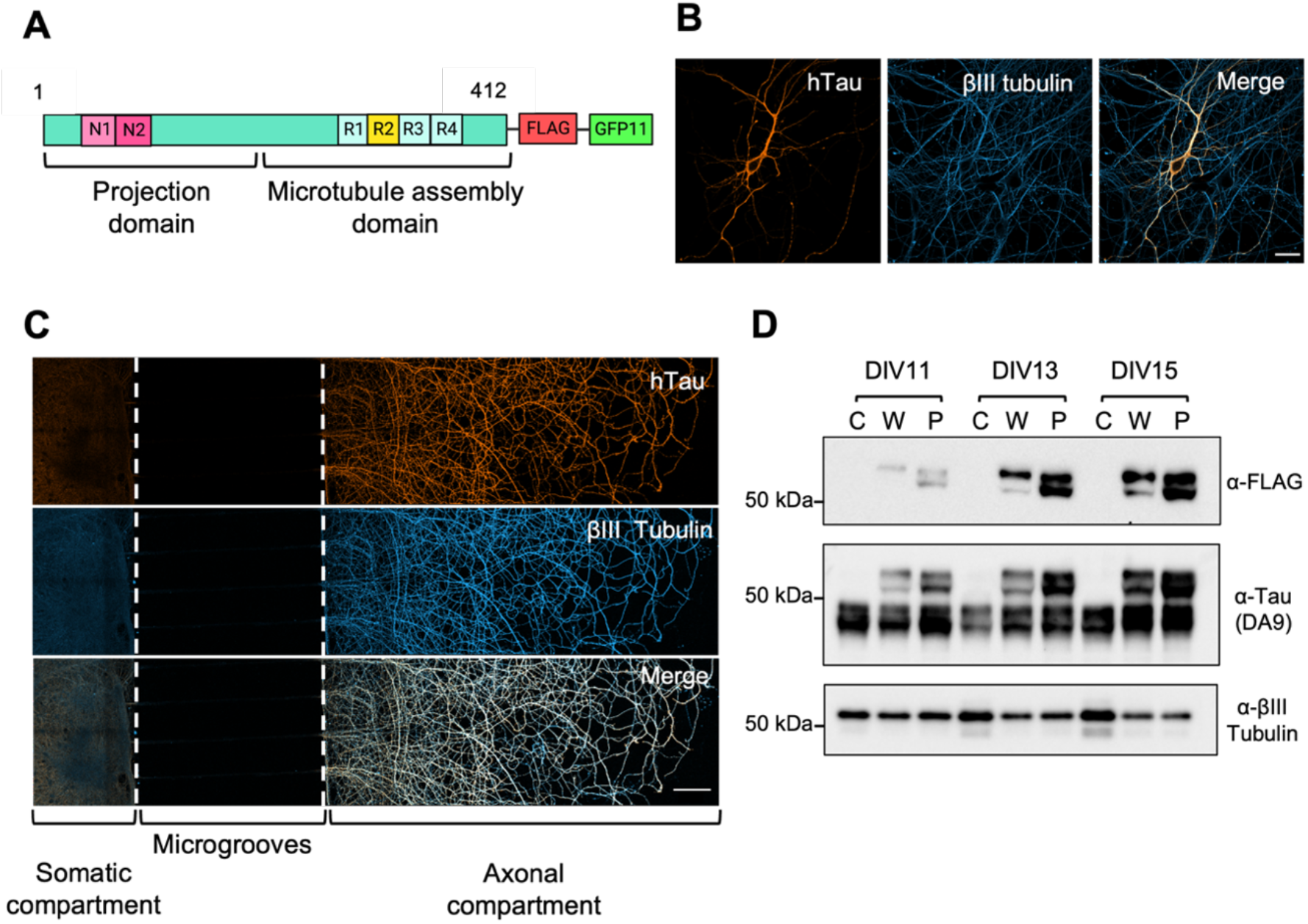
Expression of 1N4R human tau in primary hippocampal neurons. A) Schematic of tagged human tau (hTau) encoded by the pHR-hSYN lentiviral vector. The 1N4R hTau isoform (wild-type or harbouring the P301S mutation) carries a FLAG tag and a GFP11 fragment for easier detection. B-C) Primary hippocampal neurons express wild-type hTau (orange, anti-FLAG antibody), which co-localises with βIII tubulin (blue). B) DIV12 hippocampal neurons in mass culture, transfected with WT 1N4R hTau at DIV7. Scale bar = 25 μm. C) DIV15 hippocampal neurons in MFCs, transduced at DIV1 with a lentivirus expressing WT 1N4R human tau. Scale bar = 100 μm. D) Expression of hTau (anti-FLAG) and total tau (DA9) in control primary hippocampal cultures (C) or those expressing either WT (W) or P301S (P) 1N4R hTau at DIV11, DIV13, DIV15. Cells were transduced with lentiviruses at DIV7. No hTau is detectable in control samples, using either anti-FLAG or DA9 antibodies. Panel A was created with BioRender.com.

We then determined how long after lentiviral transduction hTau was consistently expressed by neurons. Hence, neurons were transduced at DIV7 with the WT and P301S lentiviruses, and the expression of hTau was analysed at different time points (DIV11, DIV13, DIV15, **Figure 2D**). We found that the expression of both forms of hTau is time-dependent and shows a similar trend, demonstrating that the presence of the P301S mutation is not affecting hTau expression (**Figure 2D**). The anti-FLAG antibody specifically recognises a band at ~55 kDa corresponding to hTau, which is expressed in neurons transduced by the lentiviruses and absent in control non-transduced neurons (**Figure 2D**). The total tau antibody DA9 was used to visualise both human (~55 kDa) and mouse tau (~45 kDa). At DIV11, hTau expression is rather low, but it increases in neurons transduced with both the WT and P301S-expressing viruses at DIV13. Whereas the expression of hTau increases with time, as expected, the levels of mouse tau appear to be largely constant in all samples. To maximise hTau expression and neuronal viability, the following tau release experiments were performed in DIV15 primary neurons transduced at DIV7.

### Detection of hTau in axonal culture media of primary hippocampal neurons

To test the hypothesis that the release of hTau is dependent on the integrity of the synaptic SNARE complex, and confirm the results found in human and mouse synaptosomes (Mazzo et al., 2022), we quantified hTau present in the culture media of primary hippocampal neurons by ELISA. To this end, we optimised a sandwich ELISA, as schematised in **Figure 3A**. Mouse tau and hTau have very similar amino acid sequences (Hernandez et al., 2019), and few antibodies are able to discriminate between them. Hence, we exploited the FLAG tag present in our hTau constructs to distinguish it from endogenous mouse tau. We used a mouse anti-FLAG antibody (coating antibody) to first capture the hTau present in the media. Then, we used a rabbit anti-total tau antibody as the detection antibody (**Figure 3A**). For assay optimisation, we built a standard curve using increasing concentrations of recombinant hTau (**Supplementary Figure 2**), which was purified based on published methods (KrishnaKumar & Gupta, 2017) and diluted in culture media. Since the amount of hTau in the axonal media was very low, we used the biotin-streptavidin system to amplify the signal, while ensuring that we retain a linear response with increasing amounts of recombinant hTau (**Supplementary Figure 2**).

**Figure 3.**
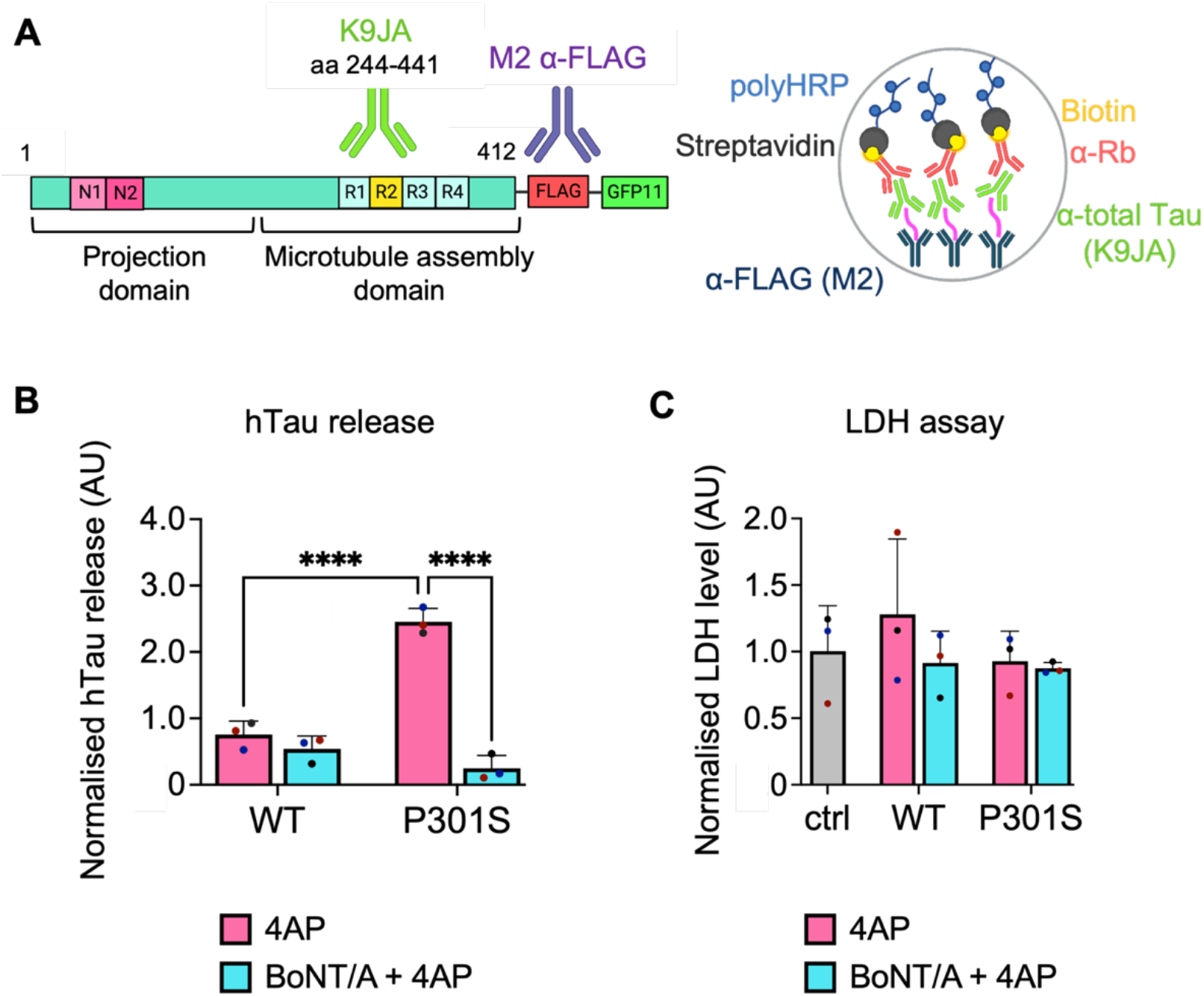
Mutant hTau release is modulated by BoNT/A. A) Schematic of the antibodies used for sandwich ELISA. A mouse anti-FLAG antibody was used to coat the plate and a rabbit total-tau antibody was used as the detection antibody. To increase the overall signal, a secondary antibody conjugated with biotin was used in combination with poly-HRP-bound streptavidin. B) Quantification of hTau release in DIV15 hippocampal neurons after treatment with BoNT/A. After 1 h treatment with BoNT/A (blue) or vehicle (magenta), samples were stimulated using 4-aminopyridine (4AP). The release of hTau is significantly higher in stimulated P301S neurons compared to either stimulated WT or P301S hTau-expressing neurons treated with BoNT/A (4AP P301S 4AP vs. WT, P301S 4AP vs. P301S BoNT/A+4AP, **** p < 0.0001). Values are normalised on the experiment average signal to account for inter-experiment variability of the ELISA. A two-way ANOVA with Bonferroni’s multiple comparisons test was performed, and all pairs were compared. n = 3. C) LDH assay of the somatic cell media upon treatment. Quantification shows absorbance at 450 nm. A one-way ANOVA with Dunnett’s multiple comparisons was used to compare treated neurons (WT, P301S) with control non-transduced cells (ctrl), and a two-way ANOVA with Šídák’s multiple comparisons test was used to compare all the treated pairs. No significant difference is detected in the LDH levels of different samples. n = 3.

### hTau release in hippocampal neurons is modulated by BoNT/A

To determine the effect of BoNT/A on hTau release in primary hippocampal neurons, we collected the media in the axonal compartments of MFCs and quantified hTau using the ELISA assay described above. Neurons were grown in the central compartment of three-chambered MFCs (**Figure 1A**, left panel) to maximise the number of axons crossing the microgrooves. Given that the volume of media in each axonal compartment is roughly half than that present in a standard two-chambered MFC (Rhymes et al., 2022), we reasoned that the concentration of released hTau in a three-chambered MFC would be approximately double, thus facilitating detection. As schematised in **Figure 1A**, neurons were transduced at DIV7. In order to prevent viruses and other molecules present in the media to diffuse from the somatic to the axonal side, the flow in the MFCs was inverted by adding a higher volume of media in the axonal compartments. For each condition analysed, we used two MFCs, for a total of four axonal compartments and two somatic compartments per data point.

At DIV15, we collected the media from all the compartments, and fresh culture media was added. Neurons were incubated for 1 h with 10 pM BoNT/A or 0.9% NaCl (vehicle) in the axonal compartment, and, after a media change, somas were stimulated with 2.5 mM 4-aminopyridine (4AP) to trigger exocytosis. 4AP inhibits voltage-gated potassium (Kv) channels, causing an intracellular increase of Ca^2+^ (Bostock et al., 1981). After 4 h of incubation, the media was collected, and neurons were fixed for immunohistochemistry.

We then quantified hTau in the media before and after treatment with the modified sandwich ELISA described in **Figure 3A** (see also Material and Methods). We found that at DIV15, when neurons are neither stimulated nor blocked, there is no difference in the release of hTau between WT and P301S-expressing neurons (**Supplementary Figure 3**). However, upon treatment, we found that P301S hTau release was significantly increased after neuronal stimulation with 4AP when compared to WT hTau (**Figure 3B**). Moreover, BoNT/A differently affected the release of two forms of hTau: the stimulated release of P301S hTau was strongly inhibited after BoNT/A treatment, whilst WT hTau release was not affected. Overall, these results suggest that the stimulated release of mutant hTau is mechanistically different from WT hTau and mediated by SNAP25-containing organelles present in the axonal terminals of neurons.

To further confirm that the differences detected in hTau release were not caused by overt membrane damage or neuronal death, we performed a lactate dehydrogenase (LDH) assay on the somatic media collected both before and after treatment with BoNT/A and 4AP (**Figure 3C**). LDH levels were identical in all experimental conditions, thus demonstrating that the differences in released hTau detected by ELISA were not driven by membrane damage. Likewise, cell viability was not affected by treatment with lentiviral particles because there was no difference in the LDH levels of transduced vs. non-transduced (control) neurons.

### hTau expression levels and axon density do not impact on hTau release

A possibility that needed to be addressed in order to offer a correct interpretation of the previous results is that the observed differences in the amount of hTau released were due to a differential expression of hTau in WT and P301S transduced neurons. To control for this variable, we measured hTau expression and axonal density in the same MFCs where we performed the release experiments. hTau expression in neurons was measured using the anti-FLAG antibody in both cell bodies and axons of hippocampal neurons (**Figure 4A-B**). No significant difference was found in hTau intensity in the soma and axons (**Figure 4B-C**) of WT and P301S neurons. Overall, these results confirm that the differences found in hTau release in **Figure 3B** were not caused by differential expression of the WT and P301S mutant, nor by differences in axonal density in MFCs.

**Figure 4.**
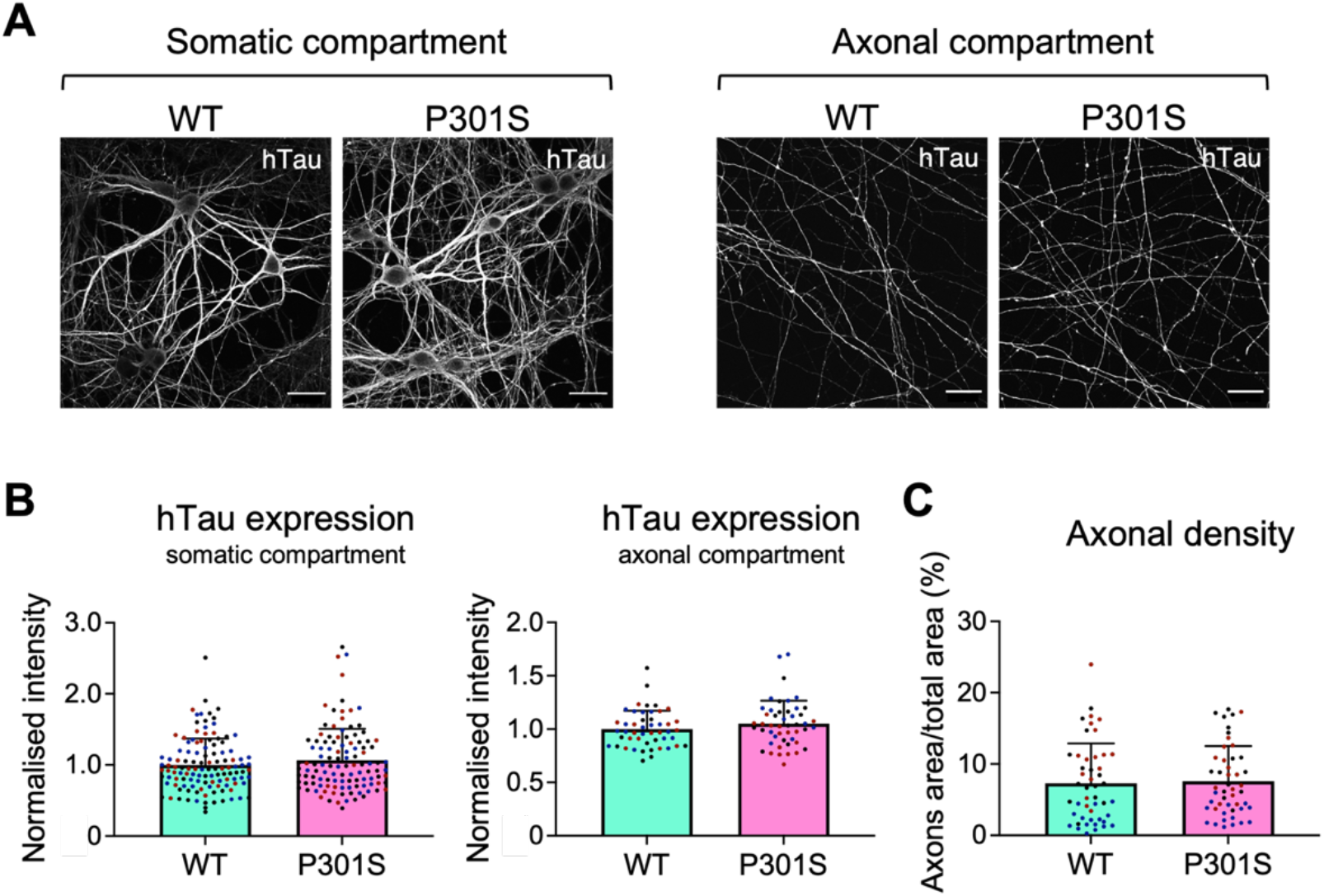
WT and P301S hTau expression levels are comparable in primary hippocampal neurons MFCs. A) DIV15 hippocampal neurons in three-compartment MFCs, transduced with WT or P301S hTau at DIV7. The expression of hTau (greyscale) in the somatic and axonal compartments was measured. Scale bar = 30 μm. B) Quantification of the signal intensity measured for WT and P301S hTau in the somatic (left panel) and axonal (right panel) chambers. For each experiment, the mean grey value was normalised by the average of the WT signal. Unpaired two-tailed Student’s t-test was used to compare WT vs P301S intensity levels in each compartment; no significant difference was found. Each dot represents a single cell body (left panel) and each field is analysed (right panel). n = 3. C) Graph showing the percentage of area occupied by axons in the axonal chambers, calculated as the axon-area/total-area ratio. Four images were taken for each MFC, and the signal intensity was measured. Unpaired two-tailed Student’s t-test was used to compare WT vs. P301S axonal density, and no significant difference was found. Single dots represent each field analysed. n = 3.

## Discussion

### BoNT/A activity in primary hippocampal neurons

BoNTs have been extensively used in both *in vitro* and *in vivo* experiments to study synaptic functionality (Caleo & Restani, 2018; Caleo et al., 2013). They target specific components of the SNARE complex, thus impairing synaptic membrane fusion and, as a consequence, neurotransmitter release. To understand whether hTau secretion was dependent on the synaptic SNARE complex in neurons, as it was in isolated synaptosomes (Mazzo et al., 2022), we decided to first use BoNT/A to target SNAP25. We used a commercial formulation of BoNT/A, abobotulinumtoxinA (Dysport, Ipsen), which is used for the treatment of several pathologies characterised by neuronal hyperactivity (Field et al., 2018). Dysport differs from other commercial preparations (Botox, Allergan and Xeomin, Merz) in its manufacturing process and the excipients and doses used in humans (Foster, 2014; Pickett, 2014). To our knowledge, the use of Dysport in primary hippocampal cultures has not been reported previously, hence we tested its activity at increasing concentrations (5-17.3 pM) in hippocampal neurons.

To detect BoNT/A activity, we used an antibody directed against its cleaved substrate SNAP25 (cl.SNAP25). Detecting cleaved substrates as a strategy to assay CNT activity enabled us to significantly amplify the signal since a single toxin molecule can cleave many SNARE molecules. This strategy has also been similarly exploited to detect BoNT/A-cleaved SNAP25 in the rodent visual system (Antonucci et al., 2008; Caleo et al., 2018; Fabris et al., 2022; Restani et al., 2011; Restani, Novelli, et al., 2012). Furthermore, this approach was shown to reliably monitor the presence of catalytic activity of BoNTs *in vitro* and *in vivo* (Antonucci et al., 2008; Restani et al., 2011).

By measuring the intensity levels of cl.SNAP25 after 24 h of incubation with BoNT/A, we found that the toxin was already active at the lower concentration used in the assay (**Figure 1B**) in unstimulated primary hippocampal neurons. The signal of cl.SNAP25 was comparable in the synapse (synaptophysin mask) and in the βIII tubulin (tubulin mask) areas in neurons (**Figure 1B**), closely matching the SNAP25 distribution in neurons (Tao-Cheng et al., 2000). Importantly, the signal obtained by antibodies directed against cl.SNAP25 in control neurons was significantly lower than in BoNT/A-treated cells (**Supplementary Figure 1**). These results demonstrate that this antibody specifically recognises cleaved SNAP25 without interacting with the full-length SNAP25 and is a reliable reporter of BoNT/A activity in intoxicated hippocampal neurons.

### P301S hTau release in primary hippocampal neurons is modulated by components of the SNARE complex

Independent lines of evidence have demonstrated the activity-dependent secretion of tau (J W Wu et al., 2013; Yamada et al., 2014) and the interaction of tau with components of SNARE complex following neuronal stimulation (Tracy et al., 2022). The SNARE complex is crucial for the fusion of neurotransmitter-containing vesicles and secretory granules with the synaptic membrane, after nerve depolarization (Han et al., 2017). This event enables the influx of calcium through calcium channels located in close proximity to active zones, which are the preferred sites of neuroexocytosis (Duman & Forte, 2003; Jahn et al., 2003). Taking these findings into consideration, we aimed to demonstrate that hTau can be released via a calciumdependent mechanism at synaptic sites in intact primary hippocampal neurons. To test this hypothesis, we used BoNT/A to selectively inactivate SNAP25 in DIV15 primary hippocampal neurons expressing WT or P301S hTau. We found no significant difference in the amount of hTau secreted by unstimulated WT and P301S neurons after 7 days of viral transduction (**Supplementary Figure 3**). However, we found that P301S hTau is efficiently secreted by hippocampal neurons upon stimulation with 4AP compared with WT hTau. 4AP represents an ideal tool for our study, since it is a potent inhibitor of voltagegated potassium (Kv) channels, and it causes an increase in intracellular Ca^2+^ and synaptic exocytosis (Bostock et al., 1981; Mihaly et al., 2001). 4AP activity can also be monitored by analysing the expression of the immediate early gene *c-fos*, which is increased upon neuronal stimulation (Bullitt, 1990; Herdegen & Leah, 1998; Krisztin-Peva et al., 2019; Toth et al., 2018; Willoughby et al., 1995). Importantly, the secretion P301S hTau was significantly decreased by treating neurons with BoNT/A, whereas the release of WT hTau remained unaffected (**Figure 4B**).

These results suggest that the secretion of P301S hTau is mediated by SNAP25 in primary neurons, as it has been shown in isolated synaptosomes from a mouse model expressing hTau bearing the P301L mutation as well as in human AD brains (Mazzo et al., 2022). Mutations in hTau trigger a change in protein folding compared to the WT protein (Lathuilière et al., 2017), which might lead to the formation of oligomers and a differential interaction with receptors and proteins present on neuronal membrane (Gonzalez-Garcia et al., 2021). Based on these findings, our working hypothesis is that, in primary hippocampal neurons, mutant P301S hTau associates with specific synaptic/axonal compartments, leading to its preferential release upon neuronal stimulation.

Overall, our novel findings could help in developing new therapeutic strategies to selectively target pathogenic tau release, thus reducing its spreading in tauopathies, without affecting the physiological role of wild-type tau.

## Materials and Methods

### Animals and tissue collection

All experiments were conducted under the guidelines of the UCL Institute of Neurology Genetic Manipulation and Ethics Committees and in accordance with the European Community Council Directive of 24 November 1986 (86/609/EEC). Animal experiments were performed under license from the UK Home Office in accordance with the Animals (Scientific Procedures) Act 1986 and were approved by the UCL Institute of Neurology Ethical Review Committee. Colonies were maintained at the Queen Square Institute of Neurology Biological Services Unit. Animals were housed in a controlled temperature and humidity environment and maintained on a 12 h light/dark cycle with access to food and water provided *ad libitum*.

### Plasmids and reagents

Chemicals were obtained from Sigma-Aldrich unless otherwise stated. Recombinant human BDNF was purchased from PeproTech (UK). Plasmids used for cloning pcDNA3-TauP301S(1N4R)-Flag-GFP11 and pcDNA3-TauWT(1N4R)-Flag-GFP11, expressing human tau (hTau), were generated by Dr Samantha De La-Rocque and were cloned into a lentiviral vector. The pHR-hSYN-EGFP used as the backbone was kindly provided by Dr Edoardo Moretto. Recombinant 1N4R hTau-Flag-GFP11 was purified based as previously described (KrishnaKumar & Gupta, 2017). Antibodies used in immunofluorescence (IF) and western blot (WB) were as follows: mouse anti-total tau (DA9, Eli Lilly, WB 1:1,000); mouse anti-FLAG (M2, WB 1:1,000, IF 1:500, ThermoFisher #F1804); chicken anti-synaptophysin (IF 1:500, Synaptic Systems #101006); chicken anti-βIII-tubulin (IF 1:500, Synaptic Systems #302-306); mouse anti-βIII-tubulin (IF 1:500, WB 1:5,000, Biolegend #801201); rabbit anti-cleaved SNAP25 (IF 1:300); rabbit anti-cleaved VAMP (IF 1:500). Antibodies against cleaved SNAP25/VAMP were a kind gift by O. Rossetto and C. Montecucco (University of Padua, Italy).

### Primary hippocampal neuron cultures

Hippocampal neurons were isolated from C57BL/6 E16-17 embryos of either sex using previously described protocols (Kaech & Banker, 2006) with minor modifications. Briefly, neurons were dissociated by incubation with Accumax (Invitrogen, #00-4666-56) and warm Hanks’ balanced salt solution (HBSS, ThermoFisher, #14175053) in a 1:1 ratio at room temperature. Cells were then resuspended in plating medium, with Minimum Essential Medium (MEM, ThermoFisher, #41090028), 10% v/v heat-inactivated horse serum, 38 mM NaCl, 0.6% v/v glucose, 1x GlutaMAX (ThermoFisher, #35050-038). Before seeding, glass coverslips and microfluidic chambers (MFCs) were prepared as previously described (Lazo & Schiavo, 2022). Cells were plated either on 13 mm glass coverslips (40,000 cells) or three-compartments MFCs (100,000 cells; **Figure 1A**) coated with 1 mg/ml poly-L-lysine (Sigma-Aldrich, #P2636) in borate buffer (0.15 M, pH 8.5). Neurons were kept in maintenance medium (Neurobasal Medium (ThermoFisher, #21103-049), 1% v/v glucose, 1% v/v penicillin/streptomycin, 1x B27, 1x Glutamax). 10 ng/ml human brain-derived neurotrophic factor (BDNF; PeproTech, UK) were added to the axonal compartment of MFCs to promote axonal growth. Primary hippocampal neurons were cultured for 12-15 days at 37°C in a 5% CO2 incubator. Half of the culture medium was replaced by fresh medium every 3-4 days.

### Lentiviral production and transduction

TauP301S(1N4R)-Flag-GFP11 and TauWT(1N4R)-Flag-GFP11 viral particles were assembled by co-transfecting hTau, packaging (PAX, Addgene plasmid #22036) and envelope plasmids (VSVG, Addgene plasmid #12259) into Lenti-X HEK293T cells with Lipofectamine 3000 (ThermoFisher, # L3000015). The medium was collected at 48 and 72 h after transfection. Medium containing lentiviral particles was concentrated using a Lenti-X concentrator (Takara Bio, Japan) and resuspended in Neurobasal (ThermoFisher). Viral particles were stored at −80°C until needed. Neurons were transduced on DIV7 by adding viral particles directly to the medium.

### Primary neuronal culture treatment

A commercial preparation of BoNT/A, (abobotulinumtoxinA, Dysport, Ipsen) was reconstituted in 0.9% sterile saline (0.9% NaCl), as indicated by the manufacturer. At DIV14, primary hippocampal neurons cultured on coverslips were treated with increasing concentrations of Dysport (5 pM, 8.7 pM, and 17.3 pM diluted in 250 μl of fresh culture medium) for 24 h, then fixed. The highest concentration (17.3 pM) was calculated to not exceed 10% of the total volume to avoid overt changes in the neuronal medium. For experiments in MFCs, culture media in the axonal compartments (**Figure 1A**, blue chambers) of hippocampal neurons at DIV15 was collected and replaced with fresh maintenance media containing either 10 pM of Dysport or 0.9% NaCl as control. Cells were incubated for 1 h at 37°C in 5% CO2; media was then removed from the chambers. Fresh maintenance medium was added and 2.5 mM of 4-aminopyridine (4AP; Sigma-Aldrich #275875) in DMSO was added in the somatic chamber (**Figure 1A**, white chamber) to stimulate neuronal activity. Neurons were kept for 4 h at 37°C in 5% CO2 before media collection and fixation.

### Enzyme-linked immunosorbent assay (ELISA)

A modified sandwich ELISA was used to detect hTau in the neuronal media. Nunc Maxisorp flat-bottomed 96-well plates (ThermoFisher) were coated with 10 μg/ml of anti-Flag M2 capture antibody in filtered PBS, and shaken for 5 min at 400-600 rpm at room temperature, before shifting them overnight at 4°C. Plates were washed three times with 0.05% PBST and blotted on paper to remove the excess of liquid before every step, unless otherwise stated. Plates were blocked with 100 μl of 5% BSA and 0.05% Tween 20 in PBS (PBST) for 1.5 h at room temperature under shaking. Purified recombinant 1N4R hTau-Flag-GFP11 was diluted in maintenance medium to build a standard curve with concentrations between 0 and 5 ng/ml. Neuronal media containing hTau was either used fresh or thawed from −80°C, spun at 10,000 *g* at 4°C for 2 min, and 100 μl of the supernatant were loaded into the wells in triplicates together with the standard curve. Samples were incubated under shaking at room temperature for 2 h. Antibodies were diluted in antibody solution (1% BSA, 0.05% PBST), added to the plate and incubated for 1 h at room temperature under shaking. Plates were consecutively incubated with: rabbit anti-total tau antibody (K9JA, Dako #A0024, 5 μg/ml), anti-rabbit secondary antibody conjugated with biotin (Invitrogen #A16114; 1:30,000), and poly-HRP-streptavidin (ThermoFisher #21140; 1:15,000). Plates were washed four times and incubated with 3,3’,5,5’-tetramethylbenzidine (TMB, 1-Step Ultra TMB solution, ThermoFisher, #34029) for 20-30 min in the dark. The reaction was stopped with 2 M sulphuric acid. The absorbance was read at 450 nm with a microplate reader (FLUOstar Omega, BMG Labtech).

### Lactate dehydrogenase (LDH) assay

Conditioned media from the somatic side of the same MFCs used for the ELISA was analysed with the Cytotoxicity LDH assay kit (Abcam #ab65393) following manufacturer’s instructions. Media samples were spun at 10,000 *g* for 2 min and 20 μl per well for each sample was added on a 96-well plate in duplicates. The absorbance was read at 450 nm with a microplate reader.

### Western blotting

Hippocampal neuron lysates were prepared in RIPA buffer (50 mM Tris-HCl pH 7.5, 150 mM NaCl, 1% NP-40, 0.5% sodium deoxycholate, 0.1% sodium dodecylsulfate, 1 mM EDTA, 1 mM EGTA) with Halt™ protease and phosphatase inhibitor cocktail (1:100, ThermoFisher #78445), and incubated on ice for 30 min. Lysates were spun at 21,000 *g* at 4°C for 15 min, and the supernatant was collected. 4x Laemmli sample buffer (15% SDS, 312.5 mM Tris-HCl pH 6.8, 50% glycerol, 10% dithiothreitol, 0.1% bromophenol blue) was added to the lysates and samples were incubated for 5 min at 95°C. Protein separation was carried out using 4-12% NuPAGE gels (ThermoFisher) and transferred onto 20% methanol-soaked polyvinylidene difluoride (PVDF) membranes (Bio-Rad, USA). Membranes were blocked in 5% BSA dissolved in Tris-buffered saline containing 0.1% Tween-20 (TBST) for 1 h at room temperature, then incubated with primary antibody diluted in 5% BSA overnight at 4°C. Membranes were washed 3×10 min in TBST, then incubated with secondary antibody (HRP-conjugated, DAKO, Denmark) diluted in 5% BSA and washed again. Immunoreactivity was detected using chemiluminescent substrates (Luminata Crescendo/Classico, Merck Millipore) and the ChemiDoc Touch Imaging System with Bio-Rad ImageLab software.

### Immunofluorescence

Coverslips or MFCs were treated with 4% paraformaldehyde and 4% sucrose in PBS for 30 min. Cells were then permeabilised in blocking solution (5% BSA, 0.1% saponin in PBS) and incubated in primary antibody (detailed above) diluted in blocking solution overnight at 4°C, washed three times in PBS, and then incubated with AlexaFluor-conjugated secondary antibodies (1:500, ThermoFisher) for 1.5 h at room temperature in the dark. Coverslips/MFCs were washed three times in PBS, then coverslips were mounted with Mowiol and MFCs with Ibidi mounting medium (ibidi #50001). Cells were imaged using a Zeiss LSM 510 or LSM 980 equipped with AiryScan2. Images were processed and analysed using ImageJ2/FIJI (version 2.9.0/1.53t). The Orange/Green/Purple colour palette was obtained from Christophe Leterrier’s GitHub repository (https://github.com/cleterrier/ChrisLUTs).

### Statistical analysis

The software GraphPad Prism 9 for MacOs (version 9.5.0 - 525) was used for all statistical analyses and to generate the plots included in the figures. Data were assumed to be normally distributed. For comparison of two groups, Student’s *t*-test was used; one- or two-way analysis of variance (ANOVA) was used when multiple groups were analysed. Levels of significance and specific tests used are indicated in figure legends. For statistical significance: *, p < 0.05, **, p < 0.01, ***, p < 0.001 and ****, p < 0.0001. Mean ± standard error of mean is shown for all plots. Independent experiments are described in this study as biological replicates. A biological replicate is defined throughout this manuscript as a set of results obtained using primary neurons isolated from mouse embryos sourced from different litter.

## Acknowledgements

We would like to dedicate this work to late Professor Matteo Caleo (University of Padova and CNR of Pisa, Italy), who was pivotal for initiating the project and for critical discussion. We thank Dr Laura Restani (CNR of Pisa, Italy) and Professor Selina Wray (Queen Square Institute of Neurology, University College London, UK) for critical discussion and experimental troubleshooting, Dr Jose Norberto S. Vargas for assistance with western blots and the personnel of the Denny Brown Laboratories (Queen Square Institute of Neurology, University College London, UK) for maintenance of mouse colonies.

## Funding

This project was supported by a Human Frontier Science Program Long-Term Fellowship LT000220/2017-L (SS), Wellcome Trust Awards 107116/Z/15/Z and 223022/Z/21/Z (GS), UK Dementia Research Institute Foundation Award UKDRI-1005 (GS), an Alzheimer’s Society PhD Studentship Grant 520 (GS) and a Medical Research Council PhD studentship (SDLR). The funders had no role in study design, data collection and analysis, decision to publish, or manuscript preparation.

## Ethic Statement

All experiments were conducted under the guidelines of the UCL Queen Square Institute of Neurology Genetic Manipulation and Ethics Committees and in accordance with the European Community Council Directive of 24 November 1986 (86/609/EEC). Animal experiments were performed under license from the UK Home Office in accordance with the Animals (Scientific Procedures) Act, 1986 and were approved by the UCL Queen Square Institute of Neurology Ethical Review Committee

## Author Contributions

Conceptualisation: CP and GS. Investigation: CP. Analysis and/or interpretation of data: CP, SS, and GS. Drafting the manuscript: CP. Revising and critical reading of the manuscript: CP, SS, SDLR, EM, OML, and GS. Approval of the final version of the manuscript: CP, SS, SDLR, EM, OML, and GS

The funders had no role in study design, data collection and analysis, decision to publish, or manuscript preparation.

## Declaration of Interests

The authors declare no personal relationships that may have influenced the work described in this manuscript.

GS was recipient of a grant (LRAP program) from Eli Lilly and Company that supported a previously published work (Mazzo et al., 2022. Metabotropic glutamate receptors modulate exocytotic tau release and propagation. J Pharmacol Exp Ther, 383:117 – 28).

## Notes

### Competing Interest Statement

The authors have declared no competing interest.

