## Supplementary Material for "Botulinum neurotoxin A modulates the axonal release of pathological tau in hippocampal neurons"

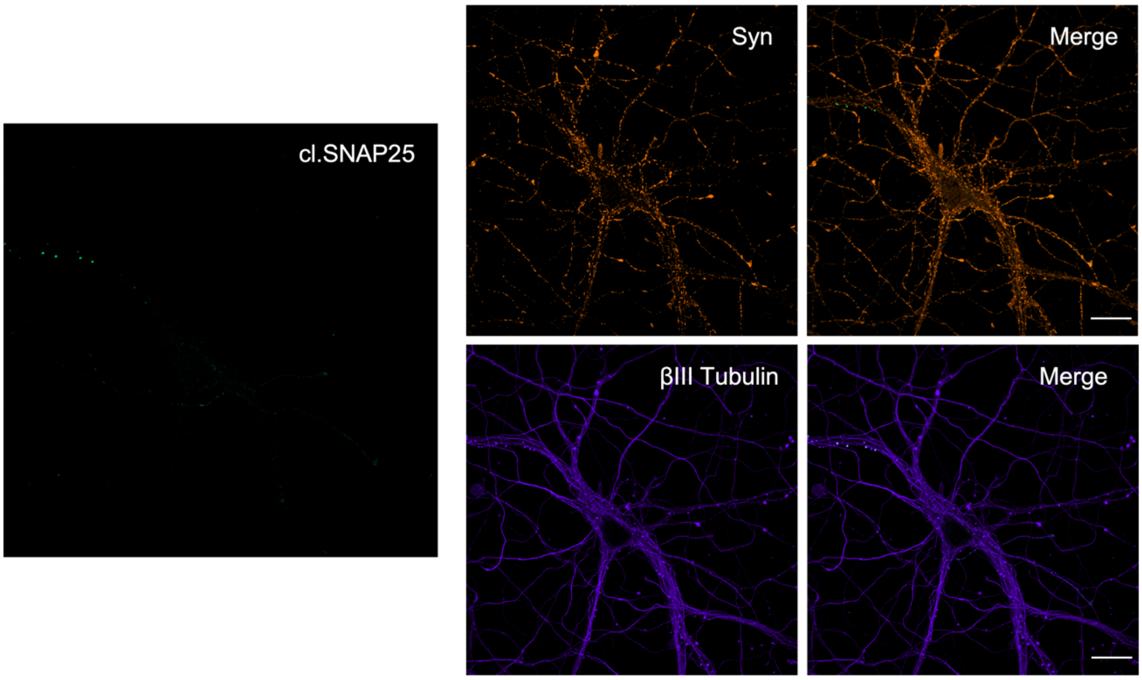


***Figure 1. The anti-cl.SNAP25 antibody gives very low or no signal in non-treated neurons.***

*Representative images of DIV15 primary hippocampal neurons showing baseline signal of cleaved SNAP25 (cl.SNAP25, green) in the absence of BoNT/A. This signal was measured in synaptophysin (Syn, orange) and βIII tubulin (purple) positive areas of non-treated neurons to determine the specificity of the antibody. No significant signal was detected. Scale bar = 25 μm.*

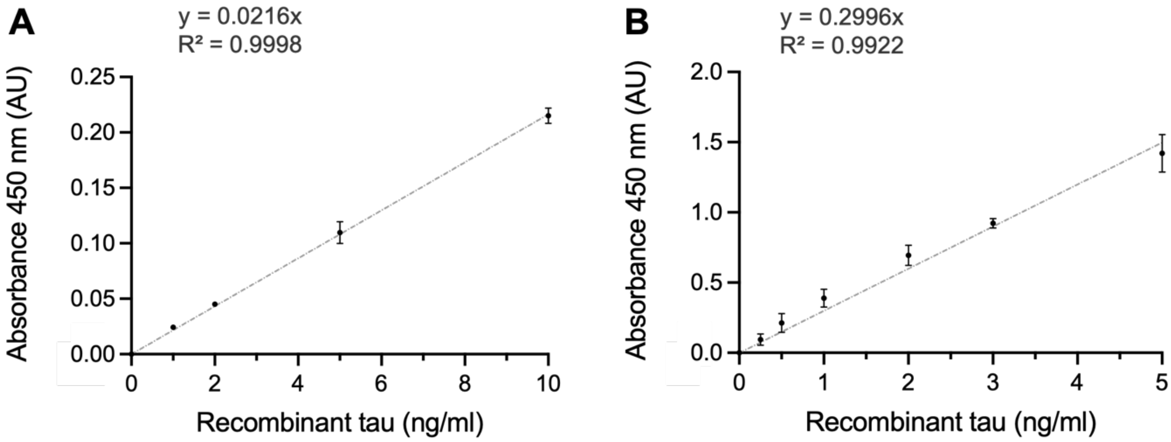


***Figure 2. Sandwich ELISA optimisation for hTau detection in neuronal media.***

*Increasing concentrations of recombinant hTau were detected using a newly developed sandwich ELISA. Graphs show the absorbance at 450 nm as a function of increasing concentrations of hTau; the intercept was set to 0. A) Absorbance at 450 nm of recombinant hTau was detected using a secondary antibody conjugated with HRP. The signal detected for the lowest concentration used (1 ng/ml) of protein is just above zero, showing the low sensitivity of this method of detection. B) Absorbance at 450 nm of recombinant hTau was detected using a biotinylated secondary antibody and then streptavidin conjugated with poly-HRP. This led to an increase in sensitivity of the assay by about tenfold, compared to A. As a result, the lowest hTau concentration detected was 0.25 ng/ml.*

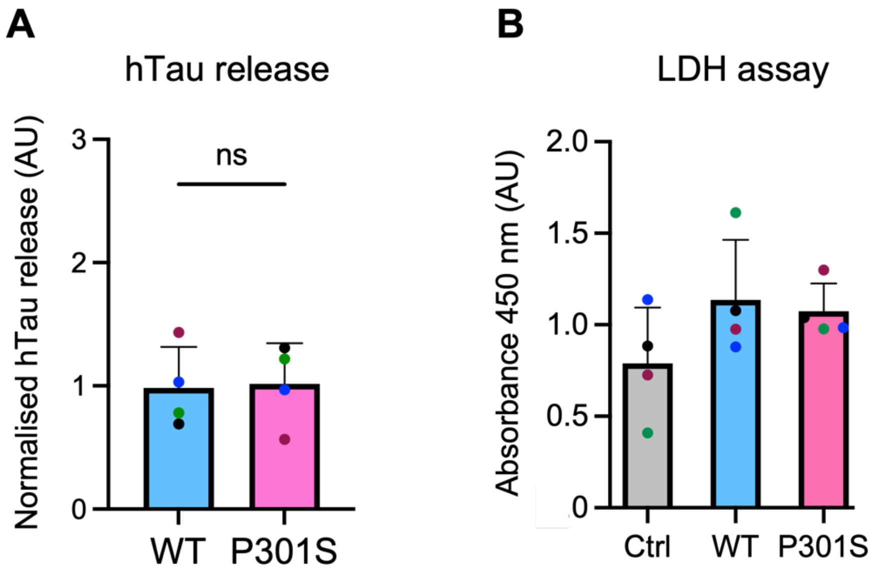


***Figure 3. Equivalent amounts of WT and P301S mutant hTau are released in the culture media of unstimulated hippocampal neurons.***

*A) Similar levels of hTau are present in the culture media of resting DIV15 primary hippocampal neurons after 7 days of lentiviral expression of WT or P301S hTau. Unpaired Student's t-test was used to compare WT and P301S, and no significant difference was detected. n = 4. B) LDH assay of the somatic cell media of DIV15 neurons. The bar chart shows absorbance levels at 450 nm. A one-way ANOVA with Tukey's multiple comparisons test was performed, and no significant difference was detected. n = 4.*
